# HLA_nc_Pred: A method for predicting promiscuous non-classical HLA binding sites

**DOI:** 10.1101/2021.12.04.471207

**Authors:** Anjali Dhall, Sumeet Patiyal, Gajendra P. S. Raghava

## Abstract

In the last two decades, ample of methods have been developed to predict the classical HLA binders in an antigen. In contrast, limited attempts have been made to develop methods for predicting binders for non-classical HLA; due to the scarcity of sufficient experimental data and lack of community interest. Of Note, non-classical HLA plays a crucial immunomodulatory role and regulates various immune responses. Recent studies revealed that non-classical HLA (HLA-E & HLA-G) based immunotherapies have many advantages over classical HLA based-immunotherapy, particularly against COVID-19. In order to facilitate the scientific community, we have developed an artificial intelligence-based method for predicting binders of non-classical HLA alleles (HLA-G and HLA-E). All the models were trained and tested on experimentally validated data obtained from the recent release of IEDB. The machine learning based-models achieved more than 0.98 AUC for HLA-G alleles on validation or independent dataset. Similarly, our models achieved the highest AUC of 0.96 and 0.88 on the validation dataset for HLA-E*01:01, HLA-E*01:03, respectively. We have summarized the models developed in the past for non-classical HLA binders and compared with the models developed in this study. Moreover, we have also predicted the non-classical HLA binders in the spike protein of different variants of virus causing COVID-19 including omicron (B.1.1.529) to facilitate the community. One of the major challenges in the field of immunotherapy is to identify the promiscuous binders or antigenic regions that can bind to a large number of HLA alleles. In order to predict the promiscuous binders for the non-classical HLA alleles, we developed a web server HLAncPred (https://webs.iiitd.edu.in/raghava/hlancpred), and a standalone package.

**Key Points:**

- Non-classical HLAs play immunomodulatory roles in the immune system.
- HLA-E restricted T-cell therapy may reduce COVID-19 associated cytokine storm.
- In silico models developed for predicting binders for HLA-G and HLA-E.
- Identification of non-classical HLA binders in strains of coronavirus
- A webserver for predicting promiscuous binders for non-classical HLA alleles

**Author’s Biography:**

1. Anjali Dhall is currently working as Ph.D. in Bioinformatics from Department of Computational Biology, Indraprastha Institute of Information Technology, New Delhi, India.
2. Sumeet Patiyal is currently working as Ph.D. in Bioinformatics from Department of Computational Biology, Indraprastha Institute of Information Technology, New Delhi, India.
3. Gajendra P. S. Raghava is currently working as Professor and Head of Department of Computational Biology, Indraprastha Institute of Information Technology, New Delhi, India.

## Introduction

Human leukocyte antigens (HLA) are an essential part of our immune system and displayed on the cell surface for antigen presentation to activate immune responses [1, 2]. In humans, the highly polymorphic HLA Class-I and II complex is located at chromosome 6p21.31 [3, 4]. More than 23000 Class-I and 8600 Class-II HLA alleles have been already reported across the globe in different ethnic groups according to IMGT/HLA database, 2020 version [5]. HLA Class-I genes further categorized into two major groups, i.e., classical (HLA-A, -B, -C) and non-classical (HLA-G, -E, -F) genes. The classical genes present antigenic peptide ligands on infected cells to CD8+ T cells and activate the immune response. On the other side, non-classical Class-I molecules moderate the immune response by inhibiting/activating stimuli of natural killer and CD8+ T cells [6]. HLA molecules protect us from several diseases by inducing and regulating immune responses [7-9]. At the same time, adverse effects have been shown in various ethnic groups due to the alteration in the expression of HLA alleles and HLA-peptide binding regions. This malfunctioning in the HLA-peptide binding grooves may lead to severe autoimmune disorders, cancer development, metastatic progression, and poor prognosis [10-13].

Recently, several studies report that the less explored non-classical alleles (HLA-G and HLA-E) play immunomodulatory roles [9, 14-16] in both innate and adaptive immune system (Figure 1). Of note, HLA-G possess four membrane bound isoforms and three soluble isoforms; interact with ILT-2, ILT-4, KIR2DL4, NKG2A/CD94 receptors, etc [17-19]. Previously, researcher believe that HLA-G molecules are only identified at the maternal-fetal interface, but recent studies revealed that the altered expression of HLA-G may leads to the development of cancers, auto-immune, inflammatory diseases and COVID-19 [20-27]. In addition, HLA-G inhibit the activation of immune cells including CD8+ T, dendritic, and natural killer cells during parasite and viral infections (such as influenza A virus, herpes, coronavirus, etc) [28-30]. These viral infections leads to the over-expression of HLA-G molecule and build an immune tolerance microenvironment. On the opposite side, HLA-E is monomorphic in nature, possesses low polymorphism and bound to the highly conserved peptides/epitopes. HLA-E regulates immune cells (NK cells and CD8+ T cells) by interacting with inhibiting receptor’s (NKG2A/CD94, NKG2B/CD94) and activating receptor (NKG2C/CD94) [31].

**Figure 1:**
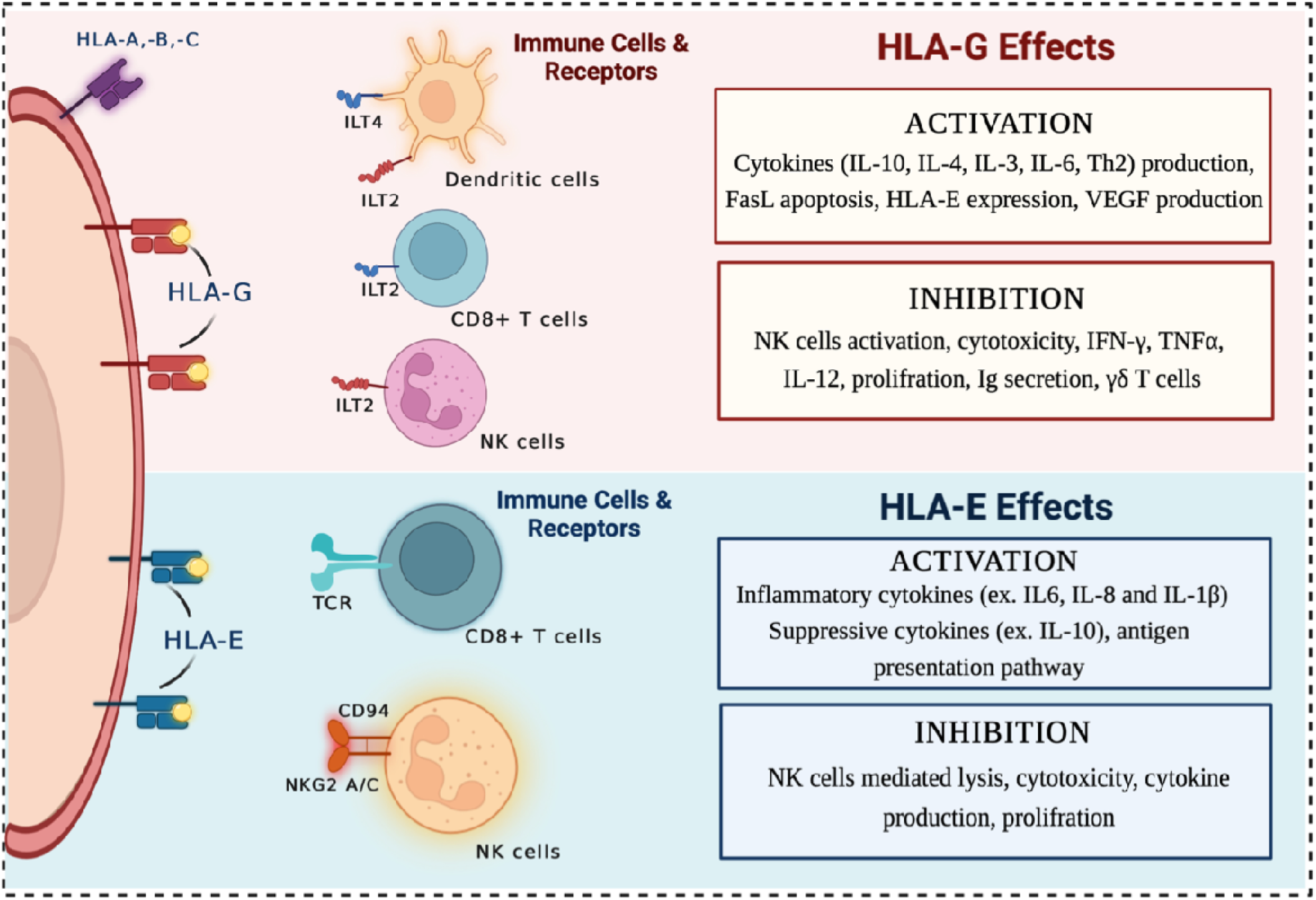
Schematic representation of non-classical HLAs (HLA-G and HLA-E) with immunomodulatory functions (inhibition/activation effects) when interact with the effective immune cells.

There are two well-known mechanisms for antigen representation by HLA-E alleles, which decide the cells’ fate. HLA-E binds to the peptide fragments descended from the signal sequence of other class Ia HLA-alleles, and this representation leads to the inhibition of NK cell functions by interacting with the NKG2A/CD94 receptors on NK cells. However, some studies revealed that the peptides from the virus (such as SARS-CoV-2, Epstein Barr Virus, Cytomegalovirus, and Hepatitis C Virus) are presented by HLA-E on the cell surface and recognised by virus specific immune cells which further activate the immune responses [32-39]. Moreover, CD8+ T-cell interaction with HLA-E leads to the production of anti-inflammatory cytokines (TGF-b, IL-4, IL-10), which is responsible for the down-regulation of pro-inflammatory cytokines production and, therefore, inhibiting the cytokine storm which plays a crucial role in the COVID-19 pathogenesis. The inhibition of cytokine storm also lowers the degree of tissue damage [40]. Several studies reveals that HLA-E inhibits NK mediated lysis, cytotoxicity, cytokine secretion, and tumor proliferation [39, 41-43], as represented in Figure 1. These findings suggest that HLA-G and HLA-E could be essential immune checkpoint molecules for designing novel immunotherapies or subunit vaccines against a wide range of diseases.

Thus, there is a need to develop methods for predicting non-classical HLA binders. Though numerous computational methods have been developed in the past for predicting HLA binder but majorly focused on classical HLA [44-50]. Only a few tools incorporate models for predicting binders for non-classical HLA alleles. Best of our knowledge no tool has been explicitly developed for predicting non-classical HLA binders. This study is dedicated for non-classical HLA, where a systematic attempt has been made to develop models for predicting non-classical HLA binders. We obtained and examined all experimentally validated non-classical HLA binders from IEDB. Based on availability of sufficient data in IEDB, we developed models for predicting binders for following non-classical alleles HLA-G*01:01, HLA-G*01:03, HLA-G*01:04, HLA-E*01:01 and HLA-E*01:03. We have used various machine learning models for the better prediction of non-classical HLA binders.

## Material and methods

### Dataset collection and pre-processing

We have collected the experimentally validated non-classical Class-I HLA-binding peptides, i.e., 1135 HLA-E and 5151 HLA-G, from the immune epitope database (IEDB), accessed on 26 October 2021. Further, we removed redundant peptides with lengths (greater than 15 or less than 8) from the dataset. After that, we obtained 142 and 723 unique peptides for HLA-E*01:01 and -E*01:03 alleles, respectively. Similarly, we get 2633, 751, and 812 unique binding peptides for HLA-G*01:01, -G*01:03, and -G*01:04 alleles, respectively. One of the major challenge in this type of study is the availability of negative dataset. Due to limited number of negative peptides in IEDB, we randomly generate the HLA-G, HLA-E non-binding peptides (length 8-15 residues) from the Swiss-Prot [51] database (March 2021 release). Finally, we obtained positive and negative dataset corresponding to each allele as represented in Table 1.

**Table 1:** Distribution of positive and negative peptides for HLA-G and HLA-E alleles obtained from IEDB and Swiss-Prot database.

| *HLA* | *HLA-allele* | *Binding Peptides*<br>*(Positive Dataset)* | *Non-Binding Peptides*<br>*(Negative Dataset)* | *Total Peptides* |
| --- | --- | --- | --- | --- |
| **HLA-G** | HLA-G*01:01 | 2633 | 2633 | 5266 |
|  | HLA-G*01:03 | 751 | 751 | 1502 |
|  | HLA-G*01:03 | 812 | 812 | 1624 |
| **HLA-E** | HLA-E*01:01 | 142 | 142 | 284 |
|  | HLA-E*01:03 | 723 | 723 | 1446 |

### Amino acid composition

Amino acid composition (AAC) of the positive and negative dataset for each allele is computed to understand the compositional similarity in different peptide sequences. The following equation, is used to compute AAC for HLA-G and HLA-E alleles binder/non-binder peptides.

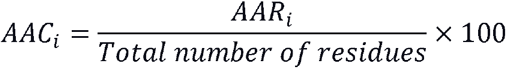

where AAC_i_ and AAR_i_ are the percentage composition and number of residues of type i in a peptide, respectively.

### Two Sample Logo (TSL)

In the current study, we performed amino-acid positional analysis using TSL tool [52], utilizing fixed-length peptide sequences used in previous studies [53-55]. In our dataset, the minimum length of peptides is eight, and hence, we create fixed-length peptides having sixteen residues. In order to create a fixed-length vector, we picked eight residues from N-terminal and eight residues from C-terminal; further, we merged the two sequences and got the final sixteen residue peptides for each positive and negative dataset.

### Feature generation

In this study, we used Pfeature [56] to calculate the binary profile for each peptide belong to binder and non-binder class. In order to calculate the binary profiles the length of the variable should be fixed, but the length of the peptides were varying from 8 to 15. Thus, to generate the fixed length vector, we extracted same number of residues from the N- and C-terminal of the peptides. Since, the minimum length of the peptides was 8, hence we chose to select the eight residues from each end and designed different patterns to calculate features. First, eight residues were selected from N-terminal, and designated as N_8_ patterns; similarly, eight residues were chosen from C-terminal and referred as C_8_ residues. Another patterns of length 16 were generated by joining the eight residues from N- and C-terminal and referred as N_8_C_8_. For the afore-mentioned patterns, each amino acid is represented by the vector of length 20, where each element represents the presence/absence of that residues, where presence was designated as “1” and absence was presented by “0”. For instance, residue “A” was represented as by vector 1,0,0,0,0,0,0,0,0,0,0,0,0,0,0,0,0,0,0,0; hence, total length of vector generated for patterns N_8_ and C_8_ was 160 (20*8) and for N_8_C_8_ its 320 (20*16). Likewise, patterns of peptides with maximum length (i.e., 15 amino acids) were generated and termed as AA_15_ and binary profile was calculated. In this case, residues “X” were added in the peptides having length less than 15. For example, 7 “X” were added in the peptide having length equal to 8, in order to make its length 15, such as peptide “VYIKHPVS” became “VYIKHPVSXXXXXXX”. In this scenario, each amino acid is represented by the vector size of 21, where last element exhibited presence/absence of “X”. Residue “V” is represented by vector 0,0,0,0,0,0,0,0,0,0,0,0,0,0,0,0,0,1,0,0,0 and “X” is presented as 0,0,0,0,0,0,0,0,0,0,0,0,0,0,0,0,0,0,0,0,1.

### Machine learning techniques

Several machine learning methods have been employed to classify the positive and negative non-classical HLA-binding peptides. Here, we have used numerous machine learning classifiers such as Decision Tree (DT), Support Vector Classifier (SVC), Random Forest (RF), XGBoost (XGB), Logistic Regression (LR), ExtraTree classifier (ET), Gaussian Naive Bayes (GNB), and k-nearest neighbors (KNN) using scikit learn python-based package [57].

### Five-fold cross-validation

We have applied a 5-fold cross-validation technique to evade the curse of biasness and overfitting in the generated models. It is one of the most crucial steps to evaluate the prediction model. In this approach, the entire dataset is partitioned into five parts, out of which four are used in training, and the resulted model is tested on the left one. The exact process is reproduced five times so that each part gets the opportunity to act as the testing dataset. Ultimately, the final performance is represented as the average of the performance of the five models that resulted from the five iterations.

### Evaluation parameters

Several parameters can be employed to assess the prediction models, which can be broadly classified into two categories termed as threshold-dependent and -independent parameters. In this study, we have determined sensitivity, specificity, accuracy, F1-score, and Matthews correlation coefficient (MCC) as the threshold-dependent parameters, whereas Area Under Receiver Operating Characteristics (AUC) curve is calculated as the threshold-independent parameter. Sensitivity (equation 1) is the measure of the ability of the model to correctly predict the binders, whereas specificity (equation 2) accounts for the percentage of correctly predicted non-binders. Accuracy (equation 3) exhibits the percentage of the correctly predicted binders and non-binders, F1-score (equation 4) captures the balance between precision and recall, and MCC (equation 5) explains the relation between the predicted and observed values. AUC is a plot between the sensitivity and 1-specificity, which captures the ability of the model to distinguish between the classes.

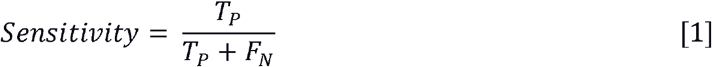

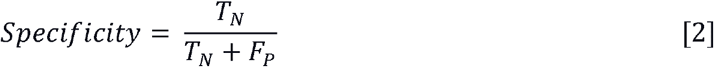

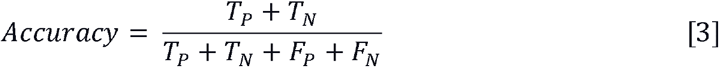

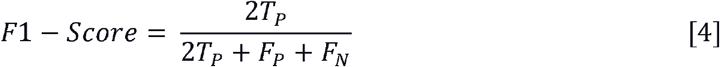

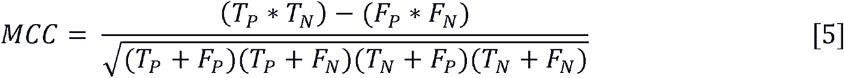

Where, T_P_, T_N_, F_P_ and F_N_ stands for true positive, true negative, false positive and false negative, respectively.

## Results

### Composition analysis

The average amino acid composition of HLA-G and HLA-E binding and non-binding peptides are represented in Figure 2. As shown in the graphs, the compositional difference in positive and negative dataset is clearly visible. HLA-G*01:01, -G*01:03, -G*01:04 alleles binders (i.e., positive peptides) have higher composition of residues such as Isoleucine (I), Lysine (K), Leucine (L) and Proline (P) as compared to non-binding peptides as depicted in Figure 2 (A, B, C). However, the average composition of Alanine (A), Leucine (L), Methionine (M), Proline (P) and Valine (V) residues is higher in HLA-E*01:01, -E*01:03 binding peptides in contrast to the negative dataset as shown in Figure 2 (D, E).

**Figure 2:**
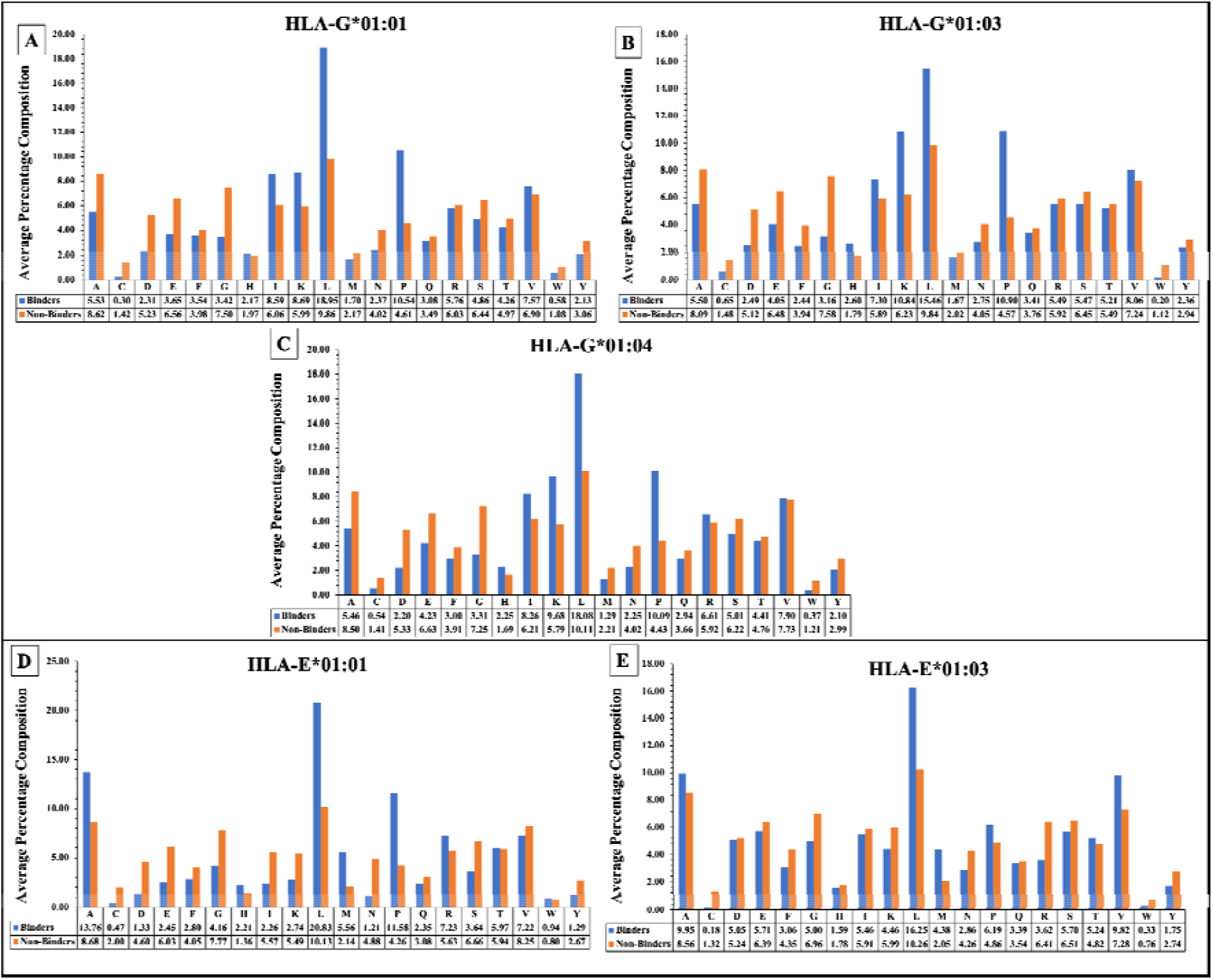
The average amino acid composition of HLA-G*01:01, -G*01:03, -G*01:04, -E*01:01, and -E*01:03 binding/non-binding peptides.

### Positional analysis

In order to analyze and study the preference or relative abundance of specific amino acid residues at a particular position in the non-classical HLA binder/non-binder sequences; we generate TSL plots (Figure 3). Of note, we generated the fixed length peptide having sixteen residues, where the first eight residues represent the N-terminal and last eight residues are C-terminal of each peptide sequence. From the TSL analysis we found that amino acid ‘L’ is predominately preferred at 2^nd^, 7^th^, 8^th^, 9^th^, 10^th^, 11^th^, and 16^th^ position in HLA-G*01:01 binding peptides (Figure 3A); at position 2^nd^, 8^th^, 9^th^, 11^th^, and 16^th^ in HLA-G*01:03 binders (Figure 3B); and at position 2^nd^, 7^th^, 8^th^, 9^th^, 11^th^, and 16^th^ in HLA-G*01:04 binding peptides (Figure 3B). Whereas, ‘I’ is most preferred at 2^nd^, 3^rd^, 7^th^, 11^th^, 15^th^, and 16^th^ position in all HLA-G alleles binders as shown in Figure 3 (A, B, C). On the other side, ‘P’ residue is preferentially located at 3^rd^, 4^th^, 7^th^, 11^th^, 14^th^, and 15^th^ positions in HLA-E*01:01 binders and 4^th^, 6^th^, 7^th^, 12^th^, and 14^th^ positions are predominantly occurred in HLA-E*01:03 binders. While ‘L’ amino acid is preferred at position 2^nd^, 8^th^, 9^th^, 10^th^ in HLA-E*01:01, however 2^nd^, 6^th^, 9^th^, 12^th^, and 16^th^ positions are mainly inhabited in HLA-E*01:03 binders.

**Figure 3:**
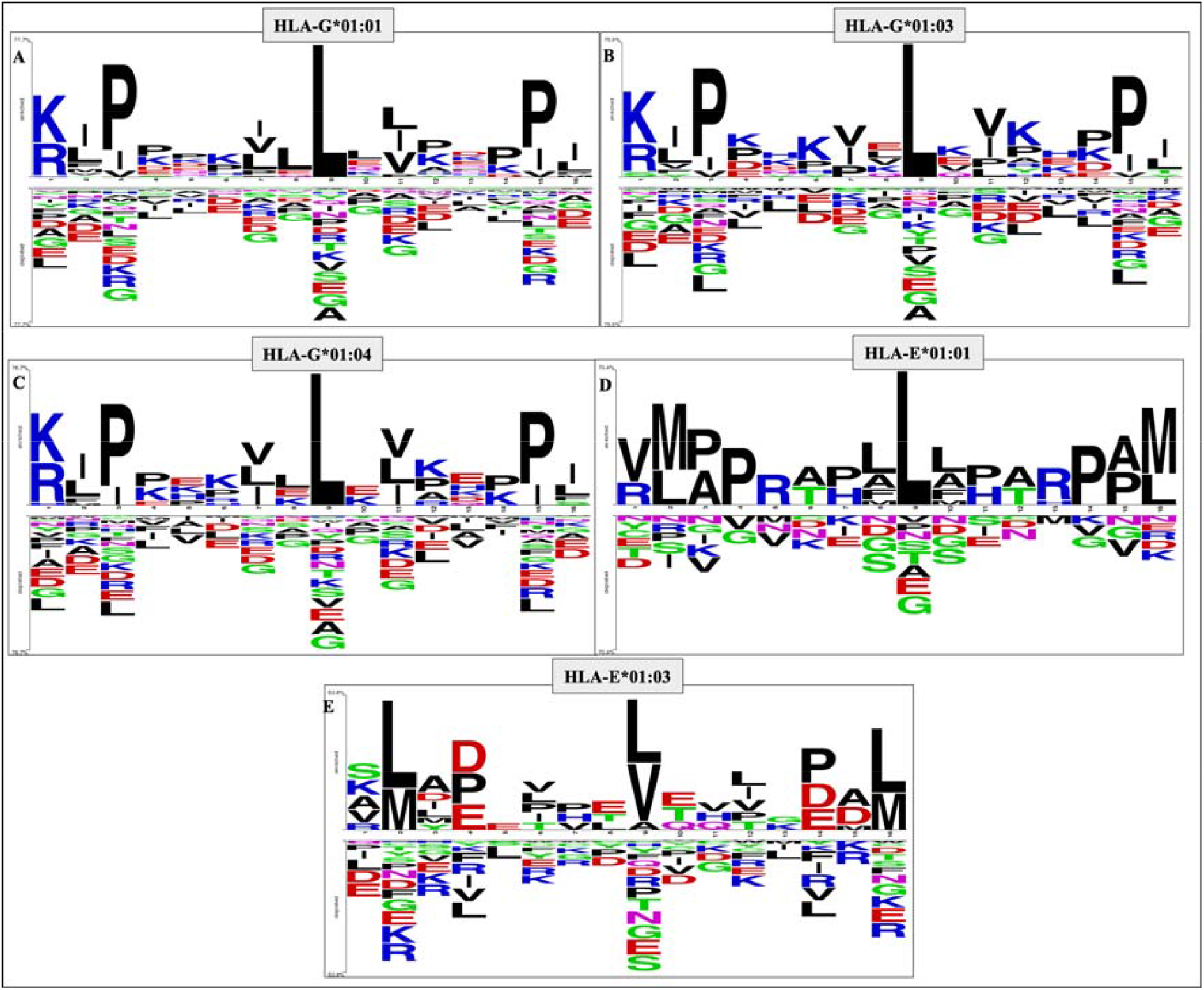
Positional abundance of amino-acid residues in HLA-G*01:01, -G*01:03, -G*01:04, -E*01:01, - E*01:03 binding/non-binding or (positive/negative) peptides, by TSL plots.

### Machine learning-based prediction

In this study, we have implemented several classifiers such as GNB, XGB, RF, DT, SVC, ET, and LR to develop prediction models. We calculate binary profile-based features of positive and negative datasets (i.e., HLA-G*01:01, -G*01:03, -G*01:04, -E*01:01, and - E*01:03 binding and non-binding peptides). Initially, we generate four feature-sets (i.e., N_8_, C_8_, N_8_C_8_, and AA_15_ binary profiles) using Pfeature standalone package. Then, we developed several machine learning models on each feature-set for HLA-G and HLA-E alleles.

### Performance of HLA-G alleles models

We compute performance of each allele dataset using various machine learning classifiers with four feature-sets as depicted in Supplementary Table S1. Further, we observed that AA_15_ binary profiles based-models outperform other feature-sets with balanced sensitivity and specificity. As shown in Table 2, HLA-G*01:01 dataset achieved maximum AUC of 0.99 and accuracy >95% on both training and validation dataset using SVC classifier (Table 2). ET based models also shows similar results on training and validation dataset with AUC of 0.99 and accuracy >95% (Table 2). XGB classifier shows comparable performance on HLA-G*01:03 dataset, with maximum AUC 0.99 and 0.98; accuracy 92.26% and 91.69% on training and validation dataset, respectively. Whereas, RF, ET, and SVC classifiers outperform the other models and perform equivalent on HLA-G*01:04 dataset. With the maximum AUC of 0.98 and accuracy 93.07% and 95.39% on training and validation dataset as shown in Table 2. On the other side, DT, LR, KNN, GBM based-models poorly performed on each dataset. The complete results for each feature-set is provided in Supplementary Table S1.

**Table 2:** The performance of machine leaning models developed using AA_15_ binary profile-based features of HLA-G alleles on training and validation datasets.

| Classifier | Training Dataset |  |  |  |  | Validation Dataset |  |  |  |  |
| --- | --- | --- | --- | --- | --- | --- | --- | --- | --- | --- |
|  | Sensitivity | Specificity | Accuracy | AUC | MCC | Sensitivity | Specificity | Accuracy | AUC | MCC |
| **HLA-G\*01:01** |  |  |  |  |  |  |  |  |  |  |
| **DT** | 88.65 | 88.03 | 88.34 | 0.93 | 0.77 | 89.37 | 89.75 | 89.56 | 0.94 | 0.79 |
| **RF** | 95.25 | 95.11 | 95.18 | 0.99 | 0.90 | 93.93 | 95.83 | 94.88 | 0.98 | 0.90 |
| **LR** | 94.26 | 94.40 | 94.33 | 0.98 | 0.89 | 92.79 | 95.07 | 93.93 | 0.98 | 0.88 |
| **XGB** | 95.44 | 95.39 | 95.42 | 0.99 | 0.91 | 94.12 | 92.98 | 93.55 | 0.98 | 0.87 |
| **KNN** | 92.93 | 92.83 | 92.88 | 0.97 | 0.86 | 91.27 | 94.12 | 92.69 | 0.97 | 0.85 |
| **GBM** | 93.31 | 79.77 | 86.54 | 0.87 | 0.74 | 91.08 | 86.34 | 88.71 | 0.90 | 0.78 |
| **ET** | 95.39 | 95.39 | 95.39 | 0.99 | 0.91 | 93.93 | 96.02 | 94.97 | 0.99 | 0.90 |
| **SVC** | 95.30 | 95.58 | 95.44 | 0.99 | 0.91 | 94.50 | 95.83 | 95.16 | 0.99 | 0.90 |
| **HLA-G\*01:03** |  |  |  |  |  |  |  |  |  |  |
| **DT** | 82.36 | 84.67 | 83.51 | 0.89 | 0.67 | 80.67 | 82.78 | 81.73 | 0.88 | 0.64 |
| **RF** | 92.18 | 92.33 | 92.26 | 0.98 | 0.85 | 87.33 | 94.04 | 90.70 | 0.97 | 0.82 |
| **LR** | 91.51 | 91.67 | 91.59 | 0.97 | 0.83 | 88.00 | 94.70 | 91.36 | 0.97 | 0.83 |
| **XGB** | 92.18 | 92.33 | 92.26 | 0.99 | 0.85 | 90.00 | 93.38 | 91.69 | 0.98 | 0.83 |
| **KNN** | 89.35 | 88.83 | 89.09 | 0.95 | 0.78 | 85.33 | 95.36 | 90.37 | 0.94 | 0.81 |
| **GBM** | 93.84 | 57.83 | 75.85 | 0.76 | 0.55 | 91.33 | 65.56 | 78.41 | 0.78 | 0.59 |
| **ET** | 92.85 | 92.67 | 92.76 | 0.98 | 0.86 | 89.33 | 94.04 | 91.69 | 0.97 | 0.84 |
| **SVC** | 92.35 | 92.50 | 92.42 | 0.98 | 0.85 | 88.00 | 95.36 | 91.69 | 0.97 | 0.84 |
| **HLA-G\*01:04** |  |  |  |  |  |  |  |  |  |  |
| **DT** | 79.54 | 80.74 | 80.14 | 0.86 | 0.60 | 86.42 | 76.69 | 81.54 | 0.87 | 0.63 |
| **RF** | 93.08 | 93.07 | 93.07 | 0.98 | 0.86 | 96.30 | 93.87 | 95.08 | 0.98 | 0.90 |
| **LR** | 92.31 | 92.30 | 92.30 | 0.98 | 0.85 | 96.30 | 92.64 | 94.46 | 0.98 | 0.89 |
| **XGB** | 92.62 | 92.76 | 92.69 | 0.98 | 0.85 | 95.06 | 94.48 | 94.77 | 0.98 | 0.90 |
| **KNN** | 89.85 | 89.99 | 89.92 | 0.96 | 0.80 | 95.06 | 90.80 | 92.92 | 0.97 | 0.86 |
| **GBM** | 90.77 | 67.49 | 79.14 | 0.79 | 0.60 | 93.21 | 67.49 | 80.31 | 0.80 | 0.63 |
| **ET** | 93.39 | 93.22 | 93.30 | 0.98 | 0.87 | 96.30 | 93.87 | 95.08 | 0.98 | 0.90 |
| **SVC** | 93.08 | 93.07 | 93.07 | 0.98 | 0.86 | 96.91 | 93.87 | 95.39 | 0.98 | 0.91 |

DT; Decision Tree, RF; Random Forest, LR; Logistic Regression, XGB; XGBoost, KNN; k-nearest neighbors, GNB; Gaussian Naive Bayes, ET; ExtraTree, SVC; Support Vector Classifier; AUC; Area Under Receiver Operating Curve, MCC; Matthews correlation coefficient.

### Performance of HLA-E alleles models

In this, we have used positive and negative datasets of HLA-E*01:01 and -E*01:03 alleles and developed various prediction models. For this dataset AA_15_ binary profile-based features prevail other classifiers as indicated in the previous section results. As demonstrate in Table 3, the performance of ET based models outperform the other classifiers with an accuracy of 87.67% and 89.47%; AUC of 0.96 on training and validation dataset, respectively. Besides, RF and XGB based models perform quite well with balanced sensitivity and specificity, accuracy (86.78% and 87.72%), AUC (0.95 and 0.96) on training and validation dataset, respectively (Table 3). However, on HLA-E*01:03 dataset RF performed quite well, with AUC of 0.93 and 0.88; accuracy of 84.78% and 82.76% on training and validation dataset as shown in Table 3. Similarly, models based on ET also perform equivalent with very less difference in sensitivity and specificity (Table 3). The comprehensive results of all other feature-sets are provided in Supplementary Table S2.

**Table 3:** The performance of machine leaning models developed using AA_15_ binary profile-based features of HLA-E alleles on training and validation datasets.

| Classifier | Training Dataset |  |  |  |  | Validation Dataset |  |  |  |  |
| --- | --- | --- | --- | --- | --- | --- | --- | --- | --- | --- |
|  | Sensitivity | Specificity | Accuracy | AUC | MCC | Sensitivity | Specificity | Accuracy | AUC | MCC |
| **HLA-E\*01:01** |  |  |  |  |  |  |  |  |  |  |
| **DT** | 78.07 | 76.99 | 77.53 | 0.82 | 0.55 | 75.00 | 75.86 | 75.44 | 0.81 | 0.51 |
| **RF** | 86.84 | 86.73 | 86.78 | 0.95 | 0.74 | 89.29 | 86.21 | 87.72 | 0.96 | 0.76 |
| **LR** | 88.60 | 89.38 | 88.99 | 0.94 | 0.78 | 85.71 | 89.66 | 87.72 | 0.97 | 0.76 |
| **XGB** | 86.84 | 86.73 | 86.78 | 0.95 | 0.74 | 89.29 | 86.21 | 87.72 | 0.96 | 0.76 |
| **KNN** | 83.33 | 83.19 | 83.26 | 0.91 | 0.67 | 82.14 | 82.76 | 82.46 | 0.93 | 0.65 |
| **GBM** | 74.56 | 76.99 | 75.77 | 0.76 | 0.52 | 89.29 | 79.31 | 84.21 | 0.84 | 0.69 |
| **ET** | 87.72 | 87.61 | 87.67 | 0.96 | 0.75 | 92.86 | 86.21 | 89.47 | 0.96 | 0.79 |
| **SVC** | 86.84 | 86.73 | 86.78 | 0.94 | 0.74 | 85.71 | 86.21 | 85.97 | 0.96 | 0.72 |
| **HLA-E\*01:03** |  |  |  |  |  |  |  |  |  |  |
| **DT** | 74.39 | 75.09 | 74.74 | 0.79 | 0.50 | 72.41 | 73.10 | 72.76 | 0.77 | 0.46 |
| **RF** | 84.95 | 84.60 | 84.78 | 0.93 | 0.70 | 82.07 | 83.45 | 82.76 | 0.88 | 0.66 |
| **LR** | 84.78 | 84.78 | 84.78 | 0.92 | 0.70 | 79.31 | 82.07 | 80.69 | 0.88 | 0.61 |
| **XGB** | 85.64 | 85.12 | 85.38 | 0.92 | 0.71 | 77.24 | 78.62 | 77.93 | 0.87 | 0.56 |
| **KNN** | 82.53 | 83.56 | 83.05 | 0.91 | 0.66 | 78.62 | 80.00 | 79.31 | 0.86 | 0.59 |
| **GBM** | 93.43 | 30.97 | 62.20 | 0.62 | 0.31 | 93.10 | 29.66 | 61.38 | 0.61 | 0.29 |
| **ET** | 84.78 | 84.60 | 84.69 | 0.93 | 0.69 | 80.00 | 84.14 | 82.07 | 0.89 | 0.64 |
| **SVC** | 85.12 | 84.95 | 85.04 | 0.93 | 0.70 | 79.31 | 76.55 | 77.93 | 0.89 | 0.56 |

DT; Decision Tree, RF; Random Forest, LR; Logistic Regression, XGB; XGBoost, KNN; k-nearest neighbors, GNB; Gaussian Naive Bayes, ET; ExtraTree, SVC; Support Vector Classifier; AUC; Area Under Receiver Operating Curve, MCC; Matthews correlation coefficient.

### Comparison with existing methods

It is very important to compare this new method with the existing methods in order to understand the advantage/disadvantages. To validate our method, we compare the performance of our models with the existing methods (MHCflurry 2.0 and NetMHCpan 4.1). We have trained our models on the previous used dataset, which is provided by MHCflurry 2.0 for HLA-G alleles only, because in case of HLA-E the number of peptides used in previous methods is very less. Further, we validate the performance of all the methods on the updated IEDB positive binders of (HLA-G*01:01, -G*01:03, and -G*01:04) alleles. As shown in the results, HLA_nc_Pred outperform other methods for the prediction of HLA-G binding peptides with balanced sensitivity, specificity, and maximum accuracy (Table 4). Whereas, MHCflurry 2.0 also performs quite well on the prediction of binder/non-binding peptides of HLA-G*01:01 dataset but perform poor on other datasets. Similarly, we observed NetMHCpan 4.1 perform poorly on the validation dataset.

**Table 4:**
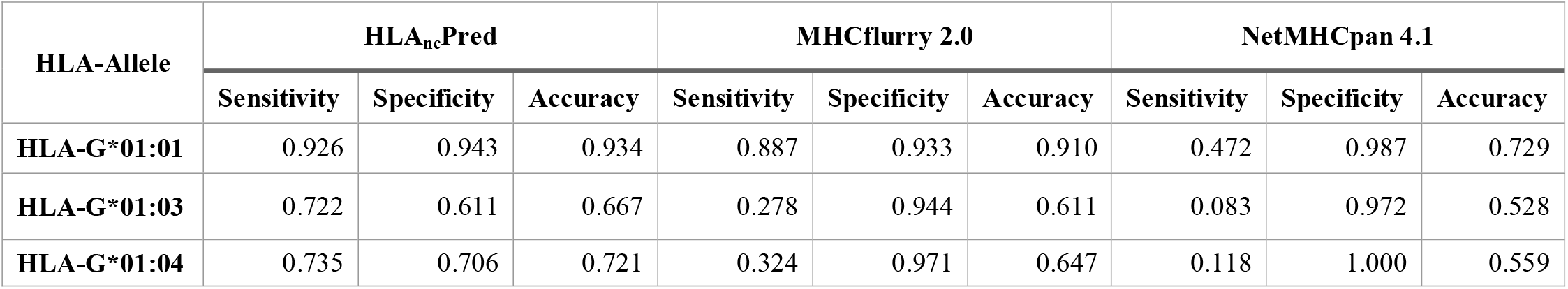
Comparison of performance of HLA_nc_Pred and other methods on the updated IEDB dataset.

| HLA-Allele | HLA<sub>nc</sub>Pred |  |  | MHCflurry 2.0 |  |  | NetMHCpan 4.1 |  |  |
| --- | --- | --- | --- | --- | --- | --- | --- | --- | --- |
|  | Sensitivity | Specificity | Accuracy | Sensitivity | Specificity | Accuracy | Sensitivity | Specificity | Accuracy |
| HLA-G*01:01 | 0.926 | 0.943 | 0.934 | 0.887 | 0.933 | 0.910 | 0.472 | 0.987 | 0.729 |
| HLA-G*01:03 | 0.722 | 0.611 | 0.667 | 0.278 | 0.944 | 0.611 | 0.083 | 0.972 | 0.528 |
| HLA-G*01:04 | 0.735 | 0.706 | 0.721 | 0.324 | 0.971 | 0.647 | 0.118 | 1.000 | 0.559 |

### Web-Server and Standalone Implementation

In the current study, for assisting the scientific community in the prediction and scanning of non-classical HLA-binder and non-binder peptides, we have developed a web-based service “HLA_nc_Pred” (https://webs.iiitd.edu.in/raghava/hlancpred/). In this web-server we have used our best models for the better prediction of HLA-binders. The detail description of HLA_nc_Pred modules are given below.

a. **PREDICT:** The prediction module allow the users to identify the most promiscuous HLA-G (-G*01:01, -G*01:02, -G*01:03) and HLA-E (-E*01:01, -E*01:03) alleles binders and non-binder peptides. User can submit multiple peptides in the standard FASTA format in the provided box or can upload the input files and can select either single or more than one alleles for binding prediction. Server provide the results in the tabular format, with the input sequence, score and prediction (binder/non-binder).
b. **SCAN:** This module gives the facility to recognize the regions of protein which may binds to the non-classical alleles such as HLA-G*01:01, -G*01:02, -G*01:03, -E*01:01, -E*01:03, using the binary profiles. It also facilitates the users to choose any length of sequence for the prediction of binders. The other way round, here users can also scan a protein sequence to find out the novel peptides with the binding ability to HLA-G and HLA-E alleles. It will generate the fragments of length selected by the users and predict their activity. User can submit the one or multiple protein sequences in FASTA format and choose the allele(s) for the prediction. In addition, user can also choose the result mode i.e., graphical or tabular as represented in Figure 4.
c. **PACKAGE:** To serve scientific community, we also provide a python- and Perl-based command line package (https://webs.iiitd.edu.in/raghava/hlancpred/stand.html) for prediction of non-classical binders at large scale and in the absence of internet. In addition, we have also develop a docker-based standalone package of HLA_nc_Pred and integrated into the ‘GPSRdocker’ package https://webs.iiitd.edu.in/gpsrdocker/ [58].
d. **DOWNLOAD:** Using this module users can download the datasets used in this study.

**Figure 4:**
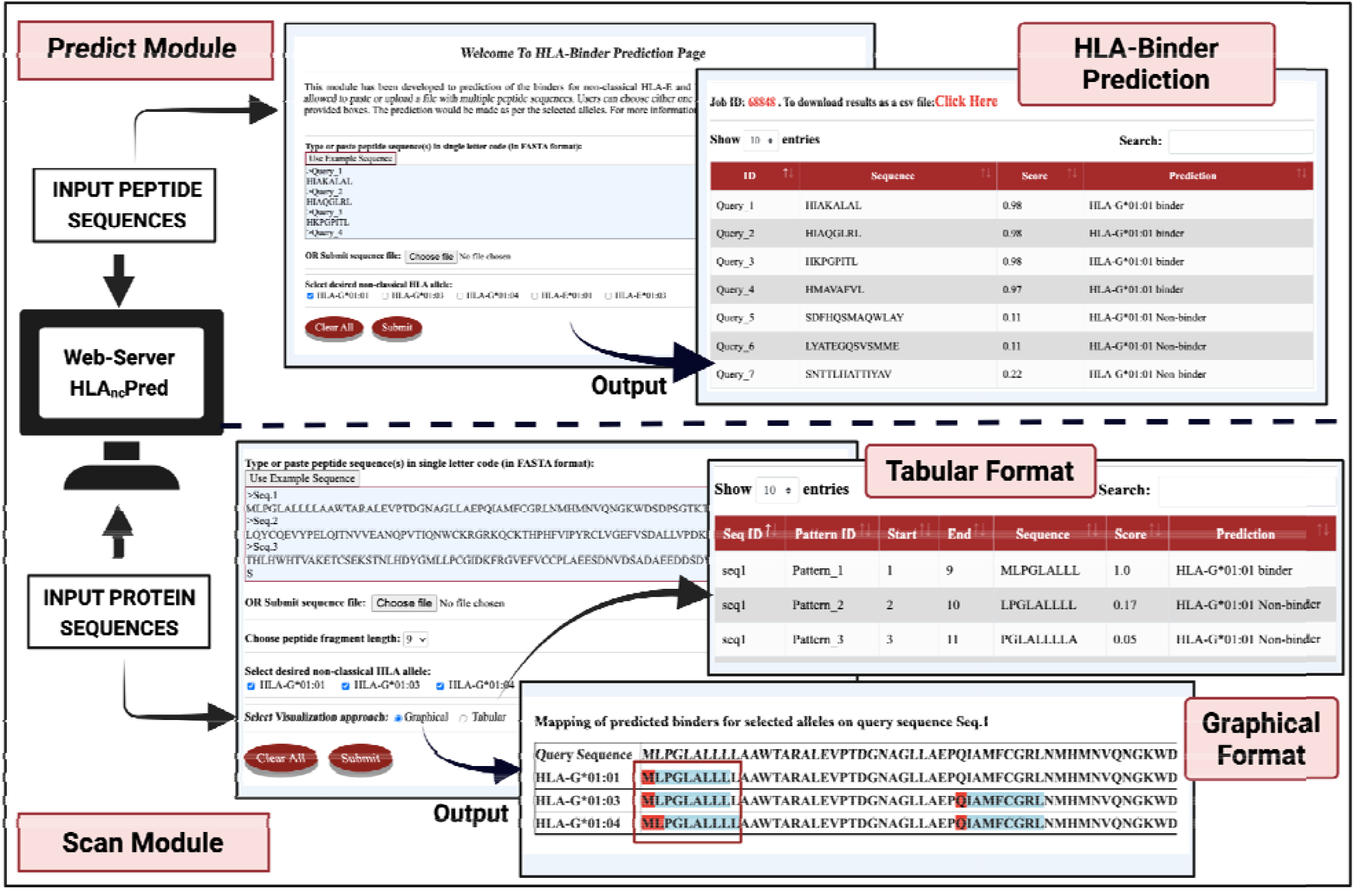
Usage of predict and scan module of HLA_nc_Pred.

The ‘HLA_nc_Pred’ web-server is compatible with a number of devices (iPhone, iPad, laptops, android mobile phones, etc.) and was build using HTML, PHP and JAVA scripts.

### Case Study: Non-classical HLA-binders in COVID-19 variants

Several studies have shown that HLA-allele binding sites significantly influence the severity of COVID-19 disease [8, 27, 40, 59]. Recently, World Health Organisation (WHO) reports that the spike protein of coronavirus has shown more than 30 mutations in the new SARS-CoV-2 variant B.1.1.529 (Omicron). In order to investigate the effect of mutations on the HLA-binding regions, we have used the reference spike protein of SARS-CoV-2 from NCBI. Further, we identified the substituted mutations named as (A67V, Δ69-70, T95I, G142D, Δ143-145, Δ211, L212I, ins214EPE, G339D, S371L, S373P, S375F, K417N, N440K, G446S, S477N, T478K, E484A, Q493K, G496S, Q498R, N501Y, Y505H, T547K, D614G, H655Y, N679K, P681H, N764K, D796Y, N856K, Q954H, N969K, L981F) associated with the spike protein of B.1.1.529 variant (https://www.gisaid.org/hcov19-variants/, https://ayassbioscience.com/voc/). In addition, we have also considered mutations in other variants such as in alpha variant (B.1.1.7) seven mutations named as N501Y, A570D, D614G, P681H, T716I, S982A and D1118H; similarly in beta variant (B.1.351) nine mutations such as D80A, D215G, K417N, E484K, N501Y, D614G, A701V, L18F and R246I; and in delta (B.1.617.2) strain nine mutations namely T19R, T95I, G142D, R158G, L452R, T478K, D614G, P681R and D950N were reported in spike protein according to Centers for Disease Control and Prevention (CDC portal) and Indian SARS-CoV-2 Genomics Consortium.

We created the mutated spike protein sequence by substituting the new mutations in the reference protein sequence. Then, we utilize the “SCAN” module (with peptide length = 9) of HLA_nc_Pred server, for the identification of HLA-binding regions in the reference and altered spike proteins. After that, we mapped the disparities in the reference and variant spike protein sequences. We found that in alpha strain, a single substitution T716I in peptide “NSIAIPINF”, results in the gain of binding ability for HLA-G*01:01; similarly E484K substitution in “KGFNCYFPL” peptide made it a binder for HLA-G*01:03, HLA-G*01:04, and HLA-E*01:01. However, due to mutation L452R in peptide “KVGGNYNYR” results in the loss of binding ability for HLA-G*01:03, and HLA-G*01:04 in delta strain. In case of recent strain omicron (B.1.1.529), mutations has resulted into the changes in the binding ability of various regions of spike proteins as shown in Table 5. The complete results for the binding prediction for alpha, beta, delta and omicron strains are provided in Supplementary Table S3, S4, S5, and S6, respectively. We hope that these findings can be used by scientific community for further investigation and designing of potential immunotherapies/vaccines against the new or future COVID-19 strains.

**Table 5:** Alterations in the binding sites of non-classical HLA alleles by mutations in Spike protein of SARS-CoV-2 variant B.1.1.529 (Omicron).

| *Alleles* | *Mutation* | *Reference Peptide* | *Mutated Peptide* | *Prediction (Binder/Non-binder)* |  |
| --- | --- | --- | --- | --- | --- |
|  |  |  |  | *(Reference)* | *(Mutated)* |
| **HLA-G\*01:01** | **Δ 211, L212I** | DLPQGFSAL | DPIGEPESA | Binder | Non-binder |
|  | **S371L** | VLYNSASFS | VLYNSASFL | Non-binder | Binder |
|  | **P681H** | PSVASQSII | HSVASQSII | Binder | Non-binder |
| **HLA-G\*01:03** | **Δ 211, L212I** | DLPQGFSAL | DPIGEPESA | Binder | Non-binder |
|  | **S371L** | VLYNSASFS | VLYNSASFL | Non-binder | Binder |
|  | **D614G** | DEVPVAIHA | GEVPVAIHA | Binder | Non-binder |
|  | **Q954H** | QNAQQLNTL | QNAQHLNTL | Non-binder | Binder |
|  | **N969K** | NSSVLNDIL | KSSVLNDIL | Non-binder | Binder |
| **HLA-G\*01:04** | **Δ 211, L212I** | DLPQGFSAL | DPIGEPESA | Binder | Non-binder |
|  | **G339D** | LCPFGEVFG | LCPFGEVFD | Non-binder | Binder |
|  | **T547K** | TLTESNKKF | KLTESNKKF | Non-binder | Binder |
|  | **N679K** | NSPRNARSV | NSPRKAHSV | Non-binder | Binder |
|  | **Q954H** | QNAQQLNTL | QNAQHLNTL | Non-binder | Binder |
|  | **N969K** | NSSVLNDIL | KSSVLNDIL | Non-binder | Binder |
| **HLA-E\*01:01** | **Δ 211, L212I** | DLPQGFSAL | DPIGEPESA | Binder | Non-binder |
|  | **S371L** | VLYNSASFS | VLYNSASFL | Non-binder | Binder |
|  | **T547K** | NGLTGTGTL | NGLTGTGKL | Non-binder | Binder |
| **HLA-E\*01:03** | **A67V, Del 69** | NVTWFHAIH | NVTWFHVIV | Non-binder | Binder |
|  | **Δ 211, L212I** | DLPQGFSAL | DPIGEPESA | Binder | Non-binder |
|  | **G339D** | LCPFGEVFG | LCPFGEVFD | Non-binder | Binder |
|  | **S371L** | VLYNSASFS | VLYNSASFL | Non-binder | Binder |
|  | **S371L, S373P, S375F** | FSTSKSYGV | FLTPKFYGV | Non-binder | Binder |
|  | **T547K** | GLTGTGTLT | GLTGTGKLT | Binder | Non-binder |
|  | **N679K** | NSPRNARSV | NSPRKAHSV | Non-binder | Binder |
|  | **D796Y** | GGDNFSQIL | GGYNFSQIL | Non-binder | Binder |

## Discussion and Conclusion

The non-classical HLA such as, HLA-G acts as an immunomodulatory molecule and natural guard while providing protection during fetus development [60, 61], whereas, HLA-E triggers immune responses via activating inflammatory cytokines during viral infections [9, 32, 62, 63]. Of note, the over-expression of HLA-G may induce the immune suppressive microenvironment, which may help in evading tumor cells from our innate and adaptive immune system. Studies have also shown that the abundant and aberrant expression of HLA-G leads to immune-mediated disorders such as multiple sclerosis, systemic lupus erythematosus [60, 64, 65]. Due to the monomorphic nature of non-classical HLA, the HLA-E restricted T cell immunotherapy may be given to a heterogenic population and possess many benefits over classical HLA-based therapies. Moreover, the activation of anti-inflammatory immune response by HLA-E interaction with CD8+ T-cell leads to the inhibition of cytokine storm and hence reduces the collateral tissue damage, which could be beneficial in treating COVID-19 patients [40]. On the other hand, HLA-G based immunotherapies have been shown to achieve encouraging results in solid-cancer treatments. Researchers have designed an anti-HLA-G CAR-T cells immunotherapy for the treatment of acute lymphoblastic leukemia and B-cell malignancies [66].

Therefore, it is the utmost need to develop an accurate prediction method for the identification of non-classical HLA-binder peptides. In last few decades, a number of HLA-peptide binding prediction methods have been developed, hitherto very limited research has been deployed in the non-classical binder prediction. To strength the previous works and to facilities the researchers working in this area, we developed a highly accurate and efficient tool dedicated to the binder prediction of non-classical HLA alleles. The dataset takes an important role in machine learning model development; thus, we have constructed our major dataset from IEDB. For the training and validation dataset we have used experimentally validated positive peptides i.e., HLA-G and HLA-E alleles binders and negative data is randomly generated using Swiss-Prot. The composition and TSL analysis indicates that non-classical HLA-binding peptides are enriched in Proline and Leucine amino acids. Furthermore, various models were developed using N_8_, C_8_, N_8_C_8_, and AA_15_ binary profiles for each allele datasets. Our results indicate that, models developed on AA_15_ binary profile-based features achieve highest performance (i.e., AUC 0.99 and accuracy >95%) on training and validation dataset for HLA-G*01:01 allele using SVC classifier. We have used the best models for each non-classical HLA-binders and developed a web-server named “HLA_nc_Pred” to predict and scan non-classical HLA-binding peptides. We also provide the standalone package for the bulk scale non-classical HLA binder prediction.

Additionally, we utilized the SCAN module of our server for the prediction of HLA-binding/non-binding peptides in the spike protein of new strain of SARS-CoV-2 i.e., B.1.1.529 (Omicron). We identified 431 promiscuous binders, where 80, 155, 56, 57, and 83 binders were predicted for HLA-E*01:01,- E*01:03, -G*01:01, -G*01:03, -G*01:04 allele, respectively. From the prediction, we observed that due to the new mutations occurred in B.1.1.529 (Omicron) variant, several peptide regions have shown the opposite binding trait in comparison with the reference spike protein (Table 5). For instance, mutations Δ 211 and L212I, changed the binding behaviour of “DLPQGFSAL” peptide for all non-classical HLA alleles. These observations can benefit the scientific community for designing vaccine against the deadly virus. In addition, researches can use this tool to predict the binding peptides of non-classical HLA-alleles against different pathogenic, autoimmune and viral infections. We anticipate that this method will benefit the community working in the area of vaccine and HLA-based immunotherapy designing. The complete architecture of the HLA_nc_Pred is shown in Figure 5.

**Figure 4:**
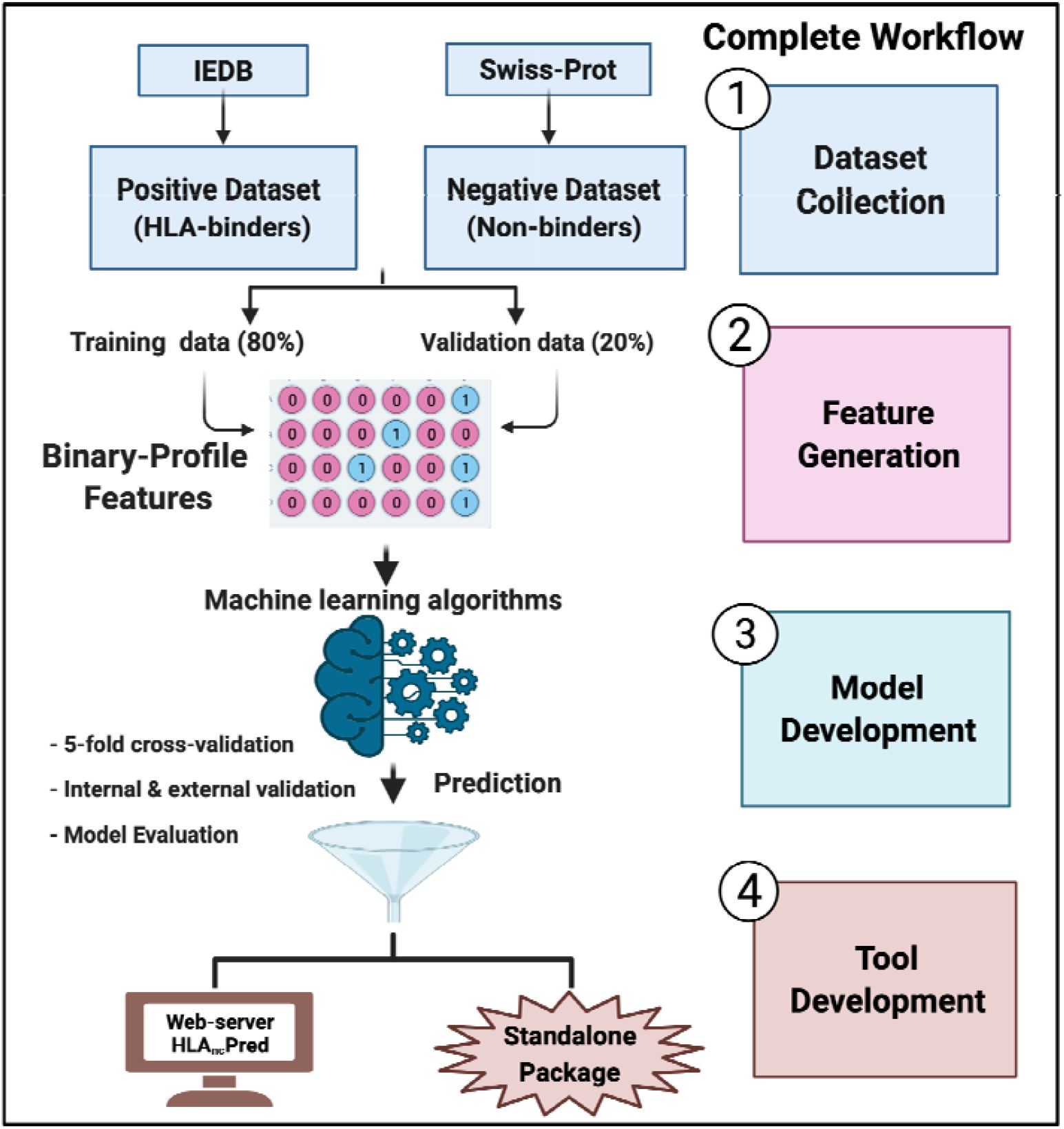
Complete architecture of HLA_nc_Pred including the dataset collection, feature generation, models and tool development.

## Authors’ contributions

AD, SP and GPSR collected and processed the datasets. AD, SP and GPSR implemented the algorithms and developed the prediction models. AD, SP and GPSR analysed the results. SP, and AD created the back-end of the web server the front-end user interface. AD, SP, and GPSR penned the manuscript. GPSR conceived and coordinated the project. All authors have read and approved the final manuscript.

## Acknowledgements

Authors are thankful to the Department of Bio-Technology (DBT) and Department of Science and Technology (DST-INSPIRE) for fellowships and the financial support and Department of Computational Biology, IIITD New Delhi for infrastructure and facilities.

## Data Availability Statement

All the datasets generated in this study are available at “HLA_nc_Pred” web server, https://webs.iiitd.edu.in/raghava/hlancpred/down.php.

